# The Potential Role of Genetic Assimilation during Maize Domestication

**DOI:** 10.1101/105940

**Authors:** Anne Lorant, Sarah Pedersen, Irene Holst, Matthew B. Hufford, Klaus Winter, Dolores Piperno, Jeffrey Ross-Ibarra

## Abstract

Domestication research has largely focused on identification of morphological and genetic differences between extant populations of crops and their wild relatives. Little attention has been paid to the potential effects of environment despite substantial known changes in climate from the time of domestication to modern day. Recent research, in which maize and teosinte (i.e., wild maize) were exposed to environments similar to the time of domestication, resulted in a plastic induction of domesticated phenotypes in teosinte and little response to environment in maize. These results suggest that early agriculturalists may have selected for genetic mechanisms that cemented domestication phenotypes initially induced by a plastic response of teosinte to environment, a process known as genetic assimilation. To better understand this phenomenon and the potential role of environment in maize domestication, we examined differential gene expression in maize (Zea mays ssp. mays) and teosinte (Zea mays ssp. parviglumis) between past and present conditions. We identified a gene set of over 2000 loci showing a change in expression across environmental conditions in teosinte and invariance in maize. In fact, overall we observed both greater plasticity in gene expression and more substantial re-wiring of expression networks in teosinte across environments when compared to maize. While these results suggest genetic assimilation played at least some role in domestication, genes showing expression patterns consistent with assimilation are not significantly enriched for previously identified domestication candidates, indicating assimilation did not have a genome-wide effect.

## Introduction

The development of agricultural societies 12,000-9,000 years ago (ka) was one of the most transformative events in human and ecological history and was made possible by plant and animal domestication^1, 2^. During domestication, crops evolved a suite of phenotypic traits, collectively known as the domestication syndrome, that distinguish them from their wild relatives^3^. Modifications due to domestication frequently include, for example, gigantism in the harvested plant part, reduced branching, and loss of shattering^3^. Scientists have sought for centuries to understand the evolution of crops during domestication, making inferences based on imperfect genetic and archaeological data. Population genetic analysis of changes associated with domestication are limited by the still sparse availability of ancient DNA, and the archaeobotanical record is often chronologically coarse and geographically uneven (e.g,^1, 2^).

As a result of these limitations, our current understanding of the morphological and molecular differences between domesticates and their wild ancestors is based almost exclusively on living representatives of those taxa.

Most of what is known about maize domestication, for example, has been drawn from comparisons between extant cultivated and wild plants. Today, profound morphological differences in vegetative architecture and inflorescence sexuality distinguish domesticated maize (*Zea mays* ssp. *mays*) and its wild ancestor teosinte (*Zea mays* ssp. *parviglumis* Iltis and Doebley; hereafter *parviglumis*). Modern teosinte has long lateral branches tipped by tassels (male inflorescences) and secondary branches bearing ears (female inflorescences) with a few small seeds covered by hard fruitcases that mature sequentially over a period of a few months. Maize, in contrast, has a single main stem terminating in a tassel and few dramatically shortened lateral branches terminated by ears instead of tassels. Maize seeds are not covered by fruitcases and its cobs mature at about the same time. These differences, the most dramatic documented for any major crop/ancestor pair, led to a century-long debate about maize ancestry^4–6^.

Because of its importance economically and as a genetic model organism, the genetics underlying the process of maize domestication has received considerable attention. Early crossing work by Beadle^4^ suggested as few as five genes could be responsible for the major vegetative architecture and inflorescence sexuality differences between maize and teosinte. More recently, work mapping quantitative trait loci (QTL) found generally consistent results, identifying six major QTL^7^. The vegetative architecture and inflorescence sexuality differences noted above, for example, are to a large degree controlled by the major QTL teosinte branched1 (*tb1*) through a change in gene expression occurring early in plant development^8–10^. Evidence of positive selection during domestication has been found at many more loci than those identified as QTL, however^11–15^, as genome-wide scans find that as much as 5% of the genome may have played a functional role in domestication^16, 17^. While there are examples such as *tga1* in which selection acted on an amino acid substitution changing the protein sequence of a gene^12^, considerable evidence suggests that much of the evolution during domestication was regulatory in nature. Not only do genes showing evidence of selection show directional changes in expression^17^, but many of the transcription and co-expression networks of maize have been substantially modified during domestication^18^ due in part to change in *cis* regulatory elements^19^.

In spite of this large body of work, domestication research has primarily focused on comparisons of extant crops and wild relatives and has largely ignored the effects of changing environmental conditions during the timeframe of crop evolution. Agricultural beginnings occurred during a period of profound global environmental change as the Pleistocene was ending and transitioning to the Holocene interglacial period^1, 20^. It is well documented that atmospheric CO_2_ and temperature were considerably lower than at present during both the Late Pleistocene (c. 14-11ka) and earliest Holocene (c. 11-9ka)^21–25^. Recent experimental work by Piperno and coauthors^26^ demonstrated remarkable phenotypic changes in teosinte exposed to temperatures and atmospheric CO_2_ similar to those experienced during the Late Pleistocene and early Holocene. These changes included maize-like vegetative architecture, inflorescence sexuality, and seed maturation, together with decreased plant height, biomass, and seed yield^26^. This work points to the possibility that early cultivators may have worked with phenotypes considerably different from those of modern teosinte. Furthermore, because some of the observed changes under experimental environments appear to have been a result of phenotypic (developmental) plasticity, the results suggest a possible role for plasticity in maize domestication^26^.

Developmental or phenotypic plasticity refers to the inherent capacity of organisms to rapidly produce novel phenotypes through one of several developmental pathways in direct response to changing environment (e.g.,^27–30^). Plasticity is now established as a mainstream concept in evolution and ecology and is increasingly considered to be fundamental for understanding the genesis of phenotypes^31–34^. Both early and recent research has also shown that genetic modifications can cement plastic phenotypes, making them stable and heritable^35, 36^. One such mechanism is genetic assimilation (GA), a process that was first investigated during the early period of the Modern Synthesis^36, 37^. Genetic assimilation involves a loss of plasticity and fixed expression across environments through reconfiguration of pre-existing genetic variation after a number of generations of growth in inducing conditions. Recent studies have demonstrated GA likely occurring in a variety of organisms, from tetrapods to *Solanum* spp. to early *Homo*, though its frequency and importance are still debated^38–40^.

Here we extend results from Piperno *et al.*^26^ on teosinte responses to environmental change, investigating the potential role of plasticity in a transcriptome-wide analysis of differential gene expression in both teosinte and maize in modern and early Holocene climate conditions. We hypothesized that expression-level changes may have constituted an initial plastic response to changing environment at the time of domestication that was later canalized through the process of GA. We find a large number of loci that show environmentally-mediated differential expression in teosinte but not maize, including some with functions consistent with phenotypic differences observed between different experimental environments and between maize and teosinte.

While population genetic evidence and enrichment analyses suggest these loci are not enriched for genes showing signals of selection during domestication, a number of loci nonetheless coincide with previously identified selective sweeps, potentially suggesting a role for GA during maize domestication. Finally, we also find a large number of genes differentially expressed in teosinte that are not identified as domestication candidates but that may nevertheless shed important light on plant responses during domestication.

## Materials and Methods

### Growth Chamber Experiment

Seeds were provided by the USDA North Central Regional Plant Introduction Station located in Ames, Iowa. We sampled four natural populations of *parviglumis* representative of the current geographic and elevational range of the subspecies^41^ as well as four maize inbred lines (Table S1).

We undertook two grow-outs in 2013 and 2014 with teosinte and maize, respectively, during their typical growing periods from July to December in two naturally-lit glass environmental chambers housed at the Gamboa field station at the Smithsonian Tropical Research Institute in Panama. One chamber was adjusted to Early Holocene (EH) temperature (ca. 23 °C) and CO_2_ (ca. 260-265 ppmv) levels determined for the low elevation Neotropics including Mesoamerica for ca. 10,000-9000 ka from paleoecological research and ice core data^21–25^. The other chamber served as a modern ambient (MA) control and was kept at ambient CO_2_ levels and temperatures, characteristic of *parviglumis* environments today^41^.

The EH chamber average CO_2_ and temperature levels were 259.7 ppmv and 23.3 °C and 260.8 ppmv and 23.2 °C in 2013 and 2014, respectively. The MA average chamber temperature was 25.1 °C in both years, with an average CO_2_ of 371 and 374 ppmv, respectively. Additional details on chamber environments can be found in^26^.

Plants were germinated from seed in five-gallon pots in natural topsoil from a local orchard and watered without fertilizer three to four times per week. In 2013 two plants were grown from each of the four *parviglumis* accessions in each chamber, followed by two replicates of each of the four maize inbreds in each chamber in 2014. We recorded weekly measurements of plant height, branch length and number, and inflorescence characteristics. After plants were harvested at maturity they were air dried and we measured the total vegetative biomass (stems, leaf, sheaths, ear bracts), node number and plant height (Tables S2 and S3).

### RNAseq Experiment

Plants were sampled for gene expression approximately 60 days after germination by removing the first visible leaf on the plant and placing it immediately in liquid nitrogen. Samples were stored at −80 °C until they were shipped overnight on dry ice to UC Davis and kept at −80 °C until extraction. Leaf tissue was ground in liquid nitrogen, and total RNA was isolated with the RNeasy mini Kit (Qiagen) following the manufacturer’s protocol. RNA quality and concentration were verified using a Bioanalyzer (Agilent RNA Nano). Total mRNA was extracted twice with Dynabeads oligo(dt)25 (Ambion) from 2*µ*g of total RNA. We prepared libraries as previously described^42^, with minor modifications and without the strand specificity. Samples were multiplexed and sequenced in two lanes of an Illumina Hiseq 2500 at the UCDavis Genome Center sequencing facility, resulting in 50 bp single-end reads with an insert size of approximately 300 bases. After demultiplexing, 3.8-20 million reads were generated for each sample (Table S4).

Low quality (base quality < 33) bases were trimmed using FASTX-Toolkit 0.0.13 http://hannonlab.cshl.edu/fastx_toolkit/ and adapters were subsequently removed using fastq-mcf version 1.04 https://code.google.com/archive/p/ea-utils/wikis/FastqMcf.wiki. Trimmed reads were mapped to the AGPv3.22 version of the maize genome using Gmap/Gsnap version 2014-05-15 with command line parameters of *-m 10 -i 2 -N 1 -w 10000 -A sam -t 8 -n 3*^43^. Read counting was performed with biocLite GenomicAlignments^44^; only reads with mapping quality 25 or higher were included in subsequent analyses. Differential gene expression was performed with DEseq2 1.10.1^45^ using a linear model (~genotype + condition) accounting for both environment (EH and MA) and population of origin; individual plants from each maize inbred line were treated as biological replicates. We used a false discovery rate (FDR) cutoff of 0.05 for determining differentially expressed genes; all results remain qualitatively identical using a more stringent cutoff of 0.01.

### Co-Expression networks

Co-expression analysis was conducted using the program WGCNA^46^. Raw expression counts were normalized using the variance stabilizing transformation in DESeq2^45^. Genes with no expression were filtered from the dataset, leaving 29,611 genes. To construct co-expression networks, Pearson correlation values were first assigned to all gene interactions and then used to create adjacency matrices by raising the correlation value to a soft power (24 and 10, for maize and teosinte, respectively). Topological overlap matrices were then formed from the adjacency matrices. The adjacency matrix indicates the connection strength between two genes (edge weights within the network), while the topological overlap matrix indicates the degree of connectivity between two genes based on their interactions with other genes in the network as well as with each other. Topological overlap matrices were used to create dissimilarity measures, which were then used to construct modules based on average linkage hierarchical clustering and the dynamic tree cut method^47^. Modules with similar eigengenes were merged using a cut-off of 0.25, meaning modules with an overall similarity of 0.75 were merged. To compare modules between EH and MA environments, a module preservation analysis was performed^48^ using EH as the reference and MA as the test for both maize and teosinte modules. Gene ontologies for each module in the maize and teosinte networks were calculated using AgriGo https://bioinfo.cau.edu.cn/agriGO/. The top hub genes were identified for each module^49^ and visualized within the module using VisANT^50^.

### Enrichment Analyses

We performed Gene ontology (GO) term enrichment analyses in AgriGo https://bioinfo.cau.edu.cn/agriGO/, using a customized reference consisting of the genes expressed in leaf tissue according to our expression data in *parviglumis* or *mays*, depending on which subspecies was used for the enrichment analysis. GO terms of all differentially expressed genes were functionally classified into three major GO categories: molecular function (MF), biological process (BP) and cellular component (CC). Genes without GO terms were removed from the analysis. We identified significantly enriched GO terms using a Fisher’s exact test and a P-value cut-off of ≤ 0.05 after applying the Yekutieli FDR correction. To test for enrichment between different categories of genes, we conducted Monte Carlo re-sampling, comparing the overlap of a particular category (e.g. teosinte-specific differentially expressed genes) with 10,000 equal-sized sets of randomly sampled genes expressed in leaf tissue.

### Additional data sets

We re-analyzed the data of Lemmon *et al.*^19^, following their methods to identify candidate genes for differential expression between maize and teosinte. For categories included in the published data (Cis only, Cis + Trans), our reanalysis identified identical gene lists. In addition to these, we followed their filtering protocol to identify a list of top candidates in Trans only and Cis x Trans regulated genes. We also used expression data from Hirsch and coauthors^51^ to calculate the coefficient of variation of expression of 48,136 genes over 503 modern inbred lines of maize to compare them to our sets of genes. Finally, we included analysis of nucleotide diversity of genes in maize and teosinte, taken from Hufford *et al.*^17^ and downloaded from https://figshare.com/articles/Gene_Popgen_Stats_from_Hufford_et_al_2012_Nat_Gen_/746968.

## Results

We grew four accessions of teosinte (*parviglumis*) and four inbred lines of domesticated maize in controlled environmental chambers simulating temperature and CO_2_ conditions reflecting Early Holocene (EH) or Modern Ambient (MA) conditions (see Methods). Many of the teosinte, particularly in the MA, had not developed inflorescences or complete branches at the time of harvest, preventing direct comparison of inflorescence sexuality. Other phenotypic characteristics we observed were nonetheless consistent with our previous experiments under these conditions^26^, with teosinte plants grown in EH conditions exhibiting smaller stature and fewer axillary nodes — indicating fewer branches — than their counterparts grown in MA (Table S2 +picture of maize/teo). Maize grown in EH conditions was also smaller and less fecund than plants in MA conditions, but in contrast to teosinte grown in previous experiments^26^ we observed no variation in branching, inflorescence sexuality, or cob development, further indicating these traits are invariant in domesticated maize (Table S3 Figure S1).

To assess differences in gene expression plasticity between teosinte and maize, we sampled leaf tissue from 39 plants and extracted and sequenced total mRNA (see Methods). On average we sampled 10 million reads per individual and identified a total of 34,341 and 35,390 expressed genes in teosinte and maize, respectively, representing 87-90 % of genes in the reference transcriptome. Analysis of differentially expressed (DE) genes under EH and MA conditions identified 3,953 and 3,355 DE genes in maize and teosinte at a false discovery rate (FDR) of 0.05 (Figure 1a; Supplemental Data File Maize DE and Teosinte DE). Many genes were differentially expressed in both taxa, and the observed 1,021 shared genes (Figure 1b) is significantly more than expected under a simple model of independence (P <1e-04). Co-expression analysis (see Methods) identified a total of 35 and 52 gene modules in maize and teosinte, respectively. Module preservation analysis indicated that gene networks were much more highly conserved between MA and EH conditions in maize than in teosinte: while only 3% of modules showed no preservation in maize, over 35% were rewired in teosinte, indicating a much more labile co-expression response of teosinte to environment (Figure 2).

**Figure 1.**
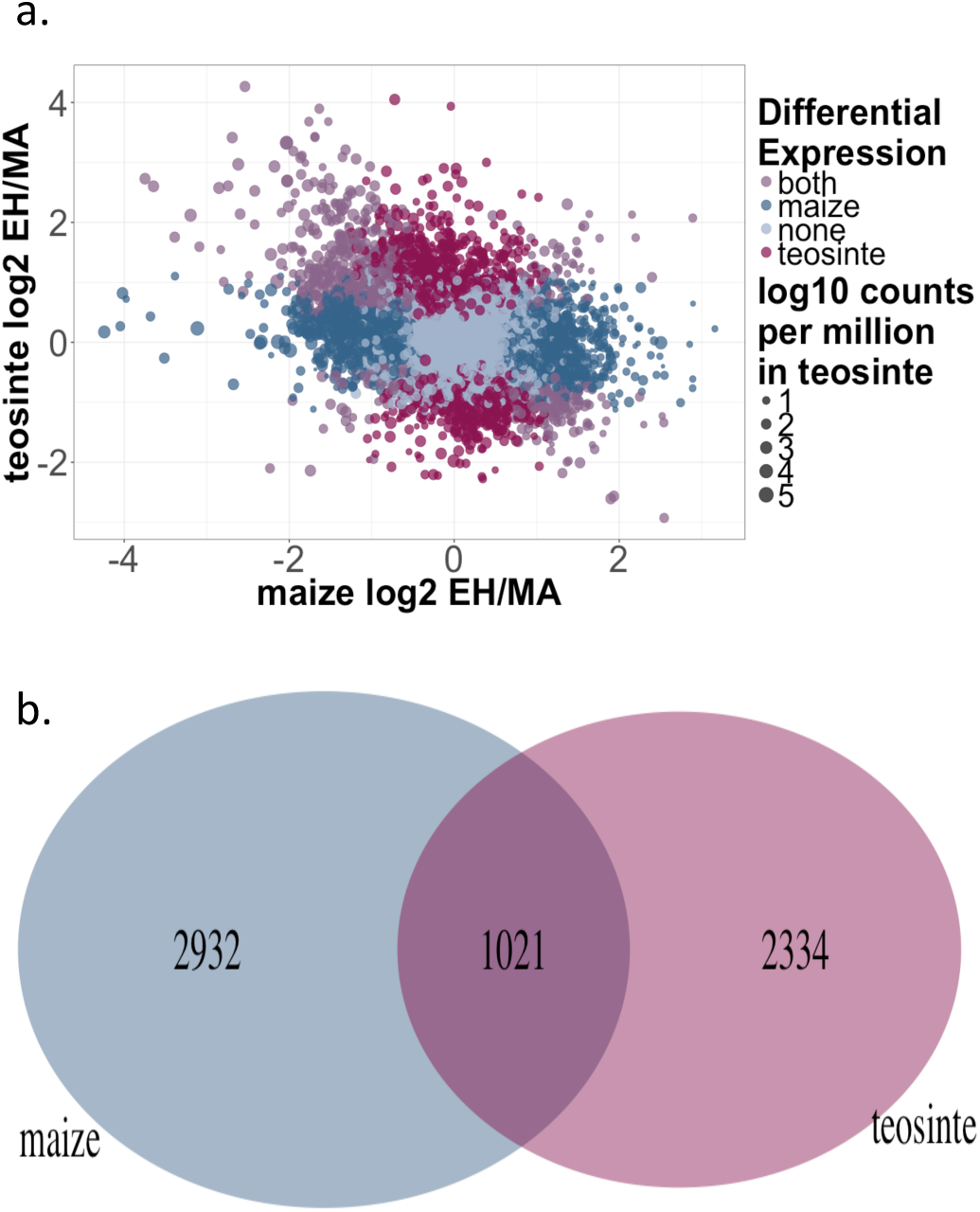
Differential expression in maize and teosinte under EH and MA conditions. (a) Categories of genes are shown in color (maize specific DE genes in blue, teosinte specific DE genes in red, shared DE genes in purple and non DE genes in gray), and point size represents the log mean counts per million in teosinte. (b) Venn diagram of the overlap (purple), among DE genes of maize (blue) and teosinte (red) when exposed to the EH environment.

**Figure 2.**
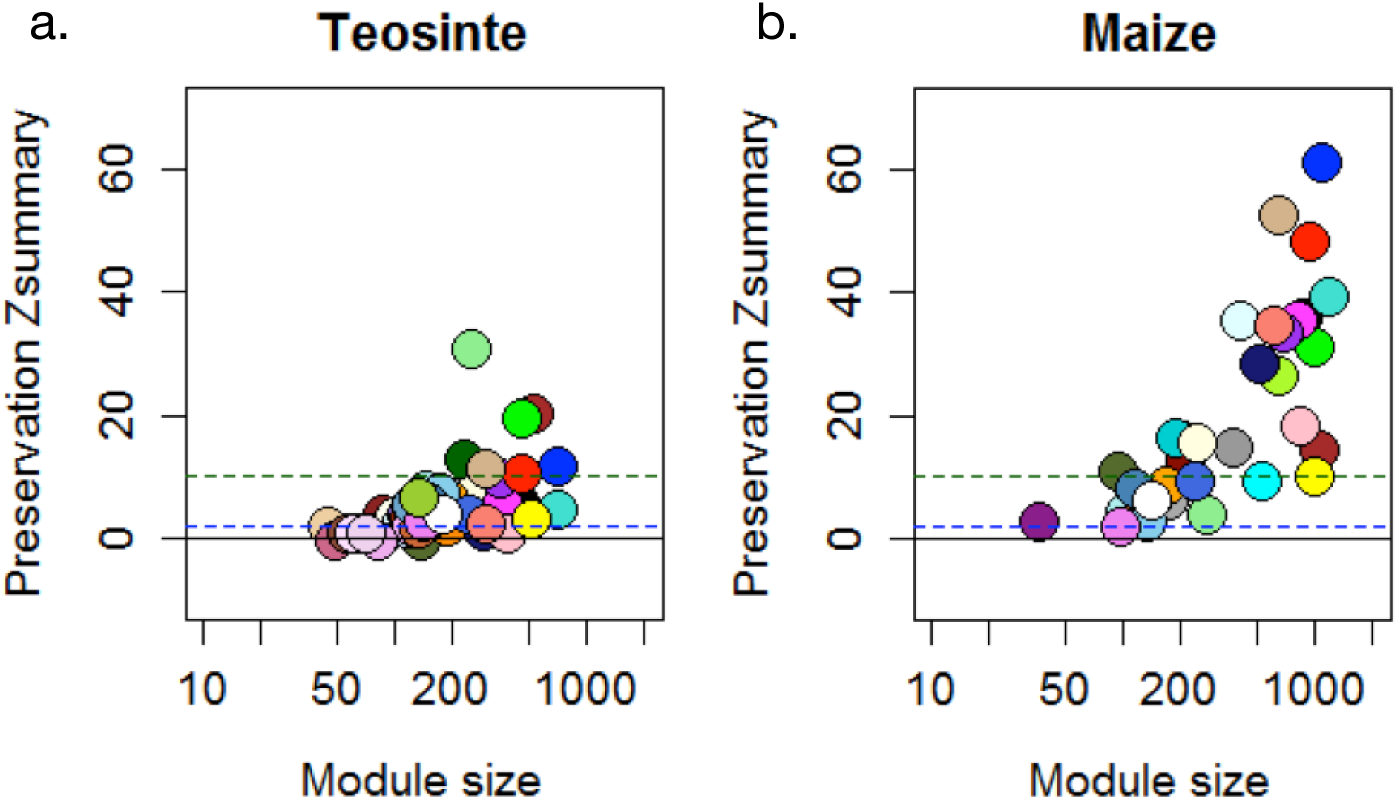
Module preservation in co-expression analysis. WGCNA preservation scores for teosinte (a) and maize (b) modules across early Holocene and modern ambient environmental conditions. Modules with scores below 2 (blue dashed line) have no preservation across conditions, those between 2 and 10 (green dashed line) are moderately preserved, and those above 10 are highly preserved

We then investigated the role of selection during domestication in shaping the observed differences in expression across environments and between teosinte and maize by taking advantage of a number of published datasets. We first reanalyzed allele-specific expression data from Lemmon *et al.*^19^ to generate lists of candidate genes with regulatory divergence between maize and teosinte (see Methods). We identified sets of genes differentially expressed in only one of the two taxa; we call these sets maize-specific and teosinte-specific DE genes. Both maize- and teosinte-specific DE gene sets were enriched for genes showing *cis* — but not *trans* — differences in expression between maize and teosinte (Figure 3). We next compared our set of taxon-specific DE genes (maize or teosinte -specific) to those showing evidence of selection during domestication^17^, but found no evidence of enrichment for candidate loci (p-value *>*0.05 in all cases; Figure 3), and maize genes exhibit similar patterns of lower nucleotide diversity when compared to teosinte across both DE and non-DE genes (Figure S3), consistent with overall patterns expected due to the demographic impacts of a domestication bottleneck^17^. Finally, we asked whether taxon-specific DE genes show different patterns of variation in expression among modern maize lines. We find that both maize- and teosinte-specific genes show reduced variation in expression across a panel of more than 500 inbred lines^51^, and teosinte-specific DE genes showed a small but statistically significant decrease in variation beyond that seen in maize-specific genes. (Figure 4).

**Figure 3.**
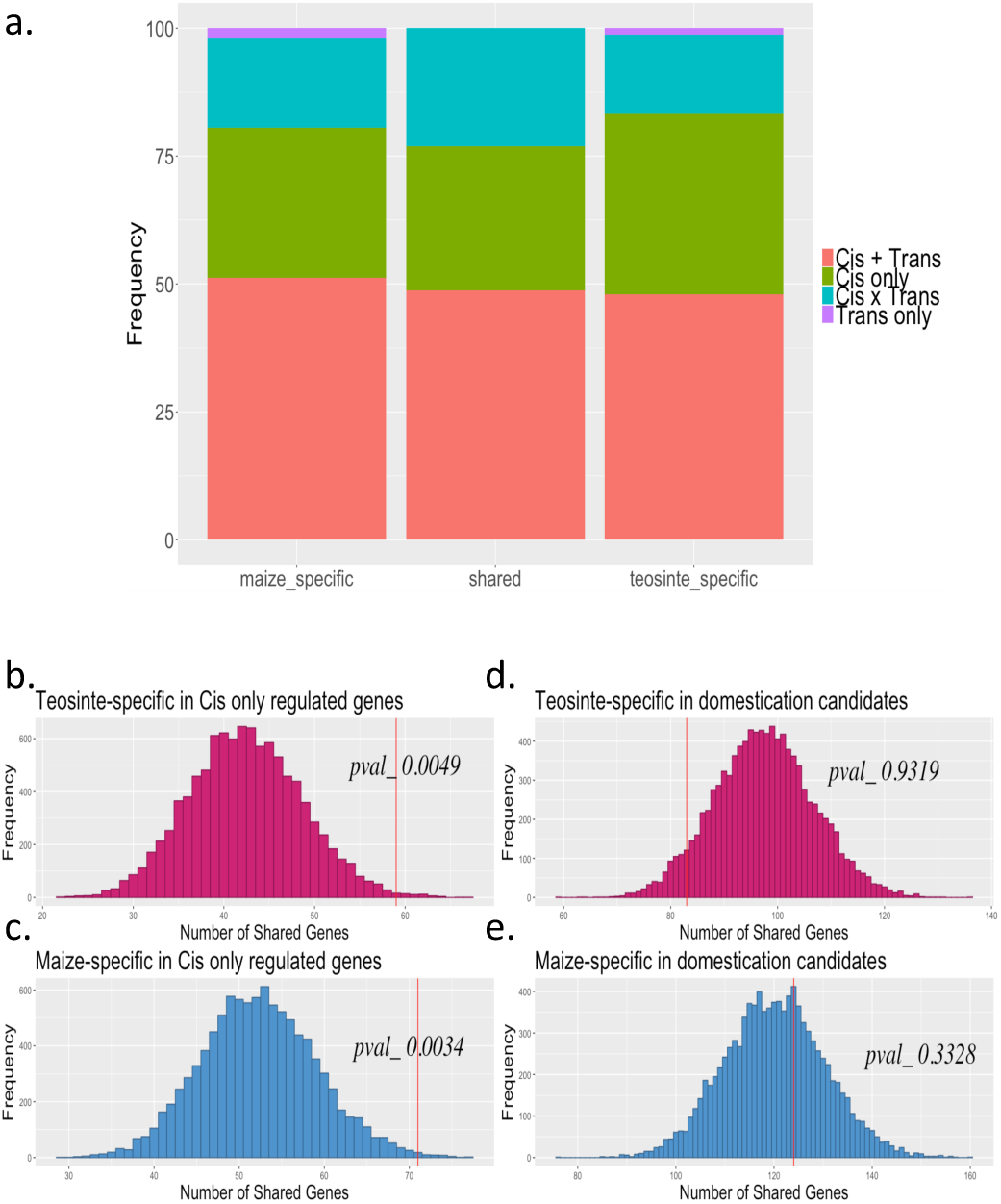
Overlap with domestication candidate genes. (a) Patterns of expression shown as a proportion of genes differentially expressed between EH and MA conditions that are also differentially expressed between maize and teosinte. Monte Carlo re-sampling of DE genes in teosinte (b,d) and maize (c,e) for enrichment in genes showing *cis*-regulated differential expression between maize and teosinte (b,c) or evidence of selection during domestication (d,e). Maize and teosinte differential expression data are from Lemmon *et al.*^19^, and selected gene lists are from Hufford et al.^17^.

**Figure 4.**
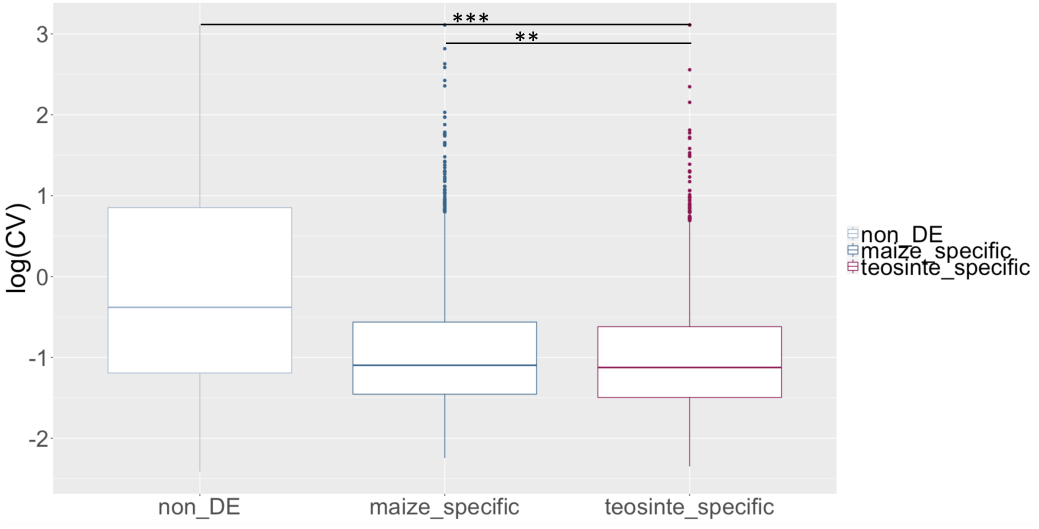
Box plot of the coefficient of variation. Genes not differentially expressed are shown in gray, maize-specific DE genes in blue, and teosinte-specific DE genes in red. The significance of the Mann-Whitney U test is as shown with **<0.01, ***<0.001

We conducted GO enrichment analysis of both shared and taxon-specific DE genes (Supplemental Data File Go terms.xls). DE genes shared between maize and teosinte are enriched in categories involved in photosynthesis, nitrogen and sugar synthesis, as well as response to stress, starvation or low phosphate conditions. Those unique to maize were mostly enriched in categories involved in photosynthesis, and these genes predominantly showed decreased expression in EH conditions; genes unique to maize also showed enrichment for biosynthesis categories. DE genes specific to teosinte were enriched for biological processes involving biosynthesis and metabolic pathways of numerous molecules including small molecules, amines, alcohols, sugars, amino acids, organic acids, and polyols. Of the few modules with co-expression rewiring in maize across environmental conditions, one module showed enrichment for ontology classes related to membrane-bounded organelles. Those rewired in teosinte showed enrichment for a diversity of ontology classes including phosphorus metabolism, protein kinase activity, organic and carboxylic acid biosynthesis, intracellular transport and localization, and amino acid ligase activity.

## Discussion

Phenotypic plasticity is a subject of growing importance in evolutionary biology^31–34^ and recent research has shown that gene expression is key to understanding both plastic and adaptive responses of plants to varying environmental conditions (e.g.,^52–54^). Several studies have shown that selection on segregating genetic variation for environmentally-induced gene expression can decrease plasticity and result in constitutive expression and even the evolution of novel traits^27, 28, 32^. This process of genetic assimilation has now been detailed in multiple taxa^38–40^ including in response to increased CO_2_^55^

In this study we sought to evaluate the role of genetic plasticity in the evolution of maize during its domestication by growing both maize and its wild ancestor teosinte in environmental conditions reflecting both modern and ancient climates. Previous experiments had demonstrated dramatic phenotypic changes in teosinte when grown under ancient conditions, and our experiment found that nearly 10% of genes expressed in leaves are differentially expressed when grown in low temperature and CO_2_ conditions reminiscent of the Early Holocene. While a similar proportion of genes also showed differential expression in maize (Figure 1), we saw much less change in overall modules of gene co-expression (Figure 2) and comparatively little change in plant morphology (Table S3).

Gene Ontology terms associated with shared and maize-specific DE genes reveal involvement in photosynthesis and are primarily down-regulated in the EH environment. Combined with GO-enrichment for stress-related genes across all candidates, these results suggest that decreases in temperature and CO_2_ were likely stressful for both maize and teosinte, and we speculate that the stress associated with ongoing rapid climate change^56^ may lead to similarly significant changes in gene expression.

While many DE genes were shared between maize and teosinte, from the perspective of domestication those showing teosinte-specific expression are of most interest, as such genes are variable in the wild ancestor but appear canalized in domesticated maize. If genetic assimilation — selection on genetic changes that canalize a plastic response such as gene expression — played a predominant role genome-wide, we might expect to see the set of teosinte-specific DE genes enriched for genes previously identified as differentially regulated between maize and teosinte^19^. While both maize- and teosinte-specific DE genes are enriched for genes showing cis-regulatory expression differences between maize and teosinte, this result is perhaps not surprising because taxon-specific DE genes were identified as genes with variable expression in one taxon and not the other. We thus expect *a priori* that these sets may have different cis-regulatory elements (and thus different response to experimental treatment) in maize and teosinte. For GA to play a genome-wide role in domestiation, we also expect genes showing evidence of canalization in maize (teosinte-specific DE genes) to show population genetic evidence of selection. Instead, we find no enrichment for genes showing evidence of selection from genome scans^17^ (Figure 3), and find that both maize- and teosinte-specific genes show decreased nucleotide diversity in maize (Figure 3), likely the result of genetic drift during the maize domestication bottleneck. While the existing evidence does not support a genome-wide impact of genetic assimilation, there are a number of reasons we might not observe such a pattern, including maladaptive plasticity^57^, selection on standing genetic variation^58^, and inbreeding during the development of modern maize lines.

Although GA may not have played a role genome-wide, our data hint at the possibility such a process may have been important for subsets of genes. For example, 83 teosinte-specific DE genes do show evidence of selection during domestication, and 6 of these have also been previously identified with a fixed regulatory difference between maize and teosinte (Supplemental Data File Teosinte specific in domestication.xls). Moreover, a number of the differentially expressed genes we observed not identified as domestication candidates have previously been linked to morphological changes similar to those important for domestication — sometimes paralleling differences between maize and teosinte — and that were previously observed in our growth chamber experiments^26,59–64^. These genes include various auxins; Brassinosteroids; a TCP transcription factor; gibberellin, absiccic acid (ABA), and cytokinin regulators; and genes implied in carbon and nitrogen fixation. Phenotypic attributes they may influence include vegetative architecture, inflorescence sexuality, plant height and biomass [e.g.,^26, 59–64^]. A relashionship between sub-optimal conditions and plasticity in teosinte is in fact well known: poor growing conditions (shade, poor soils, crowding) induce plastic phenotypic response in teosinte that include suppression of branch elongation during growth^8, 9, 65^, resulting in plants with maize attributes in vegetative and inflorescence traits similar to those seen here and in previous experiments. This suggests that these and other DE genes identified here may also lead to increased understanding of the maize domestication process by further informing the molecular basis of plasticity, phenotypic change, and adaptation in past environments. Some genes were DE only in teosinte, suggesting genetic assimilation may have occured. They include the following auxin and auxin response genes:*SAUR* 33 (GRMZM2G460861), auxin efflux carrier *PIN* 5a (GRMZM2G025742), *AUX IAA* (GRMZM2G057067) and a *PAR* (GRMZM2G423863). Also with evidence of assimilation were *TCP* (TEOSINTE-BRANCHED1/CYCLOIDEA/PCF) transcription factor 44 (GRMZM2G089361), *ZOG* 3 (GRMZM2G338465), gibberellin and ABA regulators GRMZM2G301932 and GRMZM2G338465, and nitrate reductase *NADH* 1 (GRMZM2G568636) and *ferredoxin* 1 (GRMZM2G043162).

## Conclusion

Our experimental analysis of transcriptome change has identified a large number of genes showing differential expression in maize and teosinte when grown in environments reminiscent of the Early Holocene, the time period of maize domestication. We find greater changes in teosinte morphology and gene networks, and more than 2,000 genes showing differential expression only in teosinte, suggesting substantial loss of plasticity associated with maize domestication. To our knowledge, this is the first set of transcriptomic data showing evidence of a loss of plasticity linked to domestication. Though we find little evidence to support a genome-wide role of selection and genetic assimilation in patterning this loss of plasticity, we nevertheless identify a number of genes that show evidence of genetic assimilation including some linked to morphological changes related to domestication. Future studies should expand on the work presented here by investigating additional environments (including modeled future climates) and providing more detailed, functional analysis of genes showing environmentally-induced plastic changes that may play important roles in patterning phenotypic variation in maize and teosinte.

## Acknowledgements

The authors wish to acknowledge funding support from the US National Science Foundation (IOS-0922703 and IOS-1238014) and USDA (Hatch project CA-D-PLS-2066-H) and the Smithsonian Institution. We would like to acknowledge Milton Garcia for his technical assistance. We would also like to thank members of the Ross-Ibarra lab for comments on earlier drafts of the manuscript.

## Supporting Information

**Table 1.** Sources of the teosinte and maize seeds.

| USDA accession ID | Lat/Lon | Origin |
| --- | --- | --- |
| PI 384062 | 17.417 N, -99.5 W | Valle de Bravo, Mexico State |
| PI 384063 | 18.83N, -100.16W | Mexico, Mexico State |
| PI 384071 | 18.33N, -100.31W | Iguala, Guerrero State |
| PI 566692 | 19.06N, -100.41W | Zitacuaro, Michoacán State |
| Ames 19288 Oh43 | NA | Ohio United States |
| PI 550473 B73 | NA | Iowa United States |
| NSL 30053 W22 | NA | Wisconsin United States |
| PI 558532 Mo17 | NA | Missouri United States |

**Table 2.** Teosinte phenotypes in 2013 experiment.

| All vegetative biomass g |  | Plant height cm |  | # Nodes |  |
| --- | --- | --- | --- | --- | --- |
| EHC | MCC | EHC | MCC | EHC | MCC |
| 241.2 ± 114.9 | 265.3 ± 95.8 | 141.8 ± 57.6 | 239.9 ± 48.3 | 15 ± 4.7 | 22 ± 4.5 |

**Table 3.** Maize phenotypes in 2014 experiment.

| Vegetative biomass g |  | Cobs (g) |  | Total Biomass (g) |  | Height (cm) |  |
| --- | --- | --- | --- | --- | --- | --- | --- |
| **EHC** | **MCC** | **EHC** | **MCC** | **EHC** | **MCC** | **EHC** | **MCC** |
| 55.0 ± 18.3 | 69.5 ± 17.0 | 39.3 ± 14.9 | 64.5 ± 11.7 | 94.3 ± 32.6 | 134.0 ± 25.1 | 140.6 ± 19.3 | 176.4 ± 15.5 |

**Table 4.** Number of reads per sample. Plants from a maternal source with a maize-like phenotype in a previous experiment are marked.

| Sample | Reads | Maize-like mom |
| --- | --- | --- |
| 265ppm_4B.1.txt | 3814875 |  |
| 400ppm_1A.1.txt | 4070011 | X |
| 265ppm_3C.1.txt | 4139946 | X |
| 265ppm_3A.1.txt | 4399187 |  |
| 265ppm_4C.1.txt | 4746629 | X |
| 400ppm_3C.1.txt | 5029499 | X |
| 265ppm_1B.1.txt | 5031069 |  |
| 265ppm_2B.1.txt | 5063618 | X |
| 400ppm_3B.1.txt | 5433424 |  |
| 265ppm_3B.1.txt | 5564170 |  |
| 400ppm_2B.1.txt | 5812095 | X |
| 400ppm_4A.1.txt | 5857687 |  |
| 400ppm_1A_2.1.txt | 5989185 |  |
| 265ppm_2B_2.1.txt | 6467938 |  |
| 265ppm_2A.1.txt | 6732943 |  |
| 265ppm_4A.1.txt | 7427625 |  |
| 400ppm_2B_2.1.txt | 7455893 |  |
| 400ppm_1B.1.txt | 7570271 |  |
| 265ppm_1A_2.1.txt | 7648528 |  |
| 400ppm_2A.1.txt | 7882249 |  |
| 400ppm_3A.1.txt | 8630267 |  |
| 265ppm_1A.1.txt | 8643790 | X |
| 400ppm_4C.1.tx | 8836575 | X |
| 265ppm_JRIAL9A | 14416915 |  |
| 265ppm_JRIAL9B | 13978009 |  |
| 265ppm_JRIAL9C | 15911016 |  |
| 265ppm_JRIAL9D | 14202484 |  |
| 265ppm_JRIAL9E | 16932095 |  |
| 265ppm_JRIAL9F | 15442245 |  |
| 265ppm_JRIAL9G | 14819357 |  |
| 265ppm_JRIAL9H | 14669370 |  |
| 400ppm_JRIAL9I | 14114563 |  |
| 400ppm_JRIAL9J | 14914449 |  |
| 400ppm_JRIAL9K | 20128181 |  |
| 400ppm_JRIAL9L | 18545881 |  |
| 400ppm_JRIAL9M | 16442158 |  |
| 400ppm_JRIAL9N | 15726969 |  |
| 400ppm_JRIAL9O | 16940428 |  |
| 400ppm_JRIAL9P | 17291888 |  |

**Figure 1.**
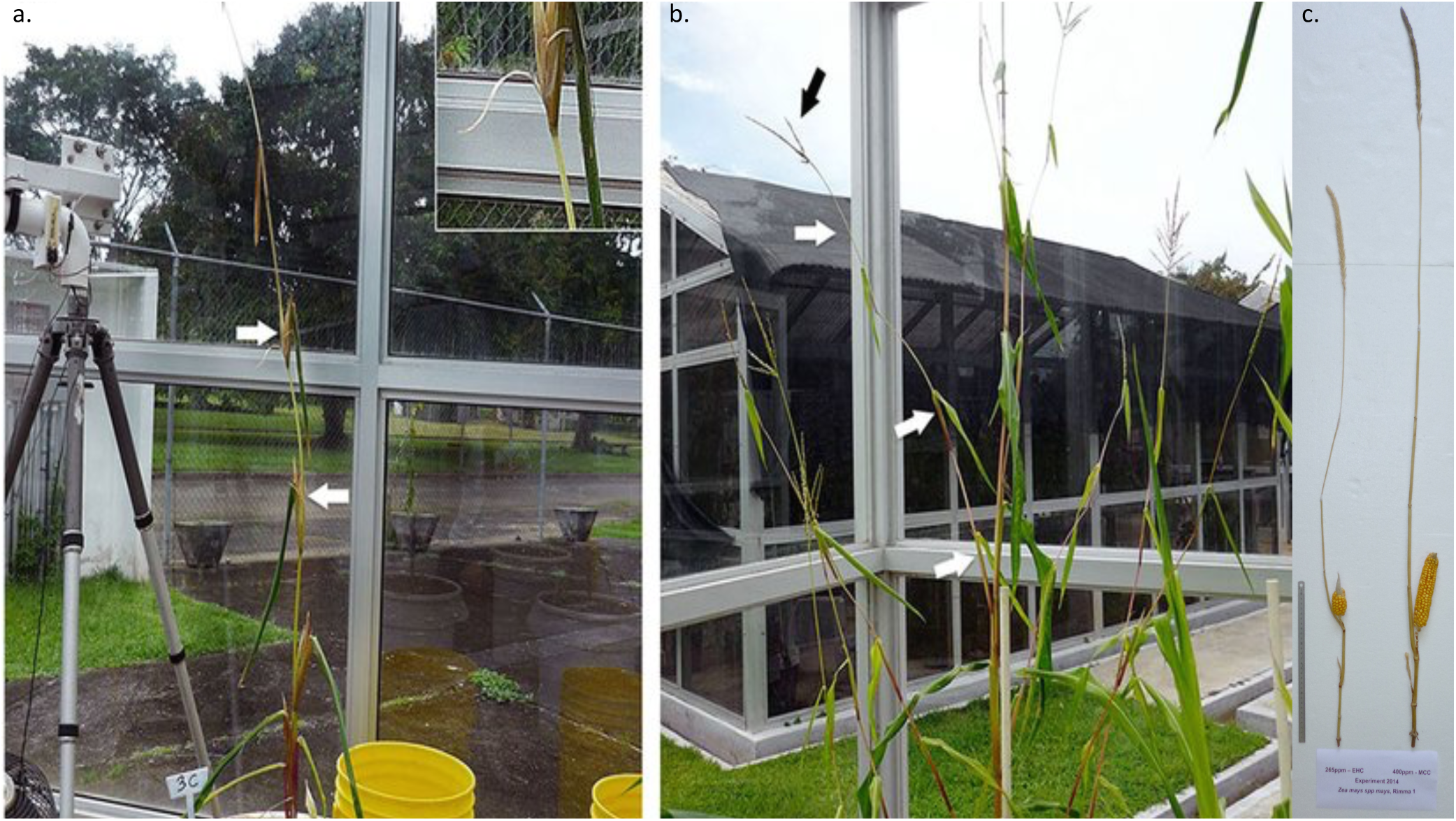
Examples of phenotypic differences in EH chamber on the left and MA chamber on the right for teosinte (a. and b. from Piperno and coauthors^26^)and maize (c.). The teosinte plant in the EH chamber is a maize-like phenotype in vegetative architecture, inflorescence sexuality, and seed maturation, as described in the main text. The plant in the MA chamber is typical of teosinte today in those characteristics. These traits are unaltered for the maize plant between the EH chamber on the left and the MA chamber on the right.

**Figure 2.**
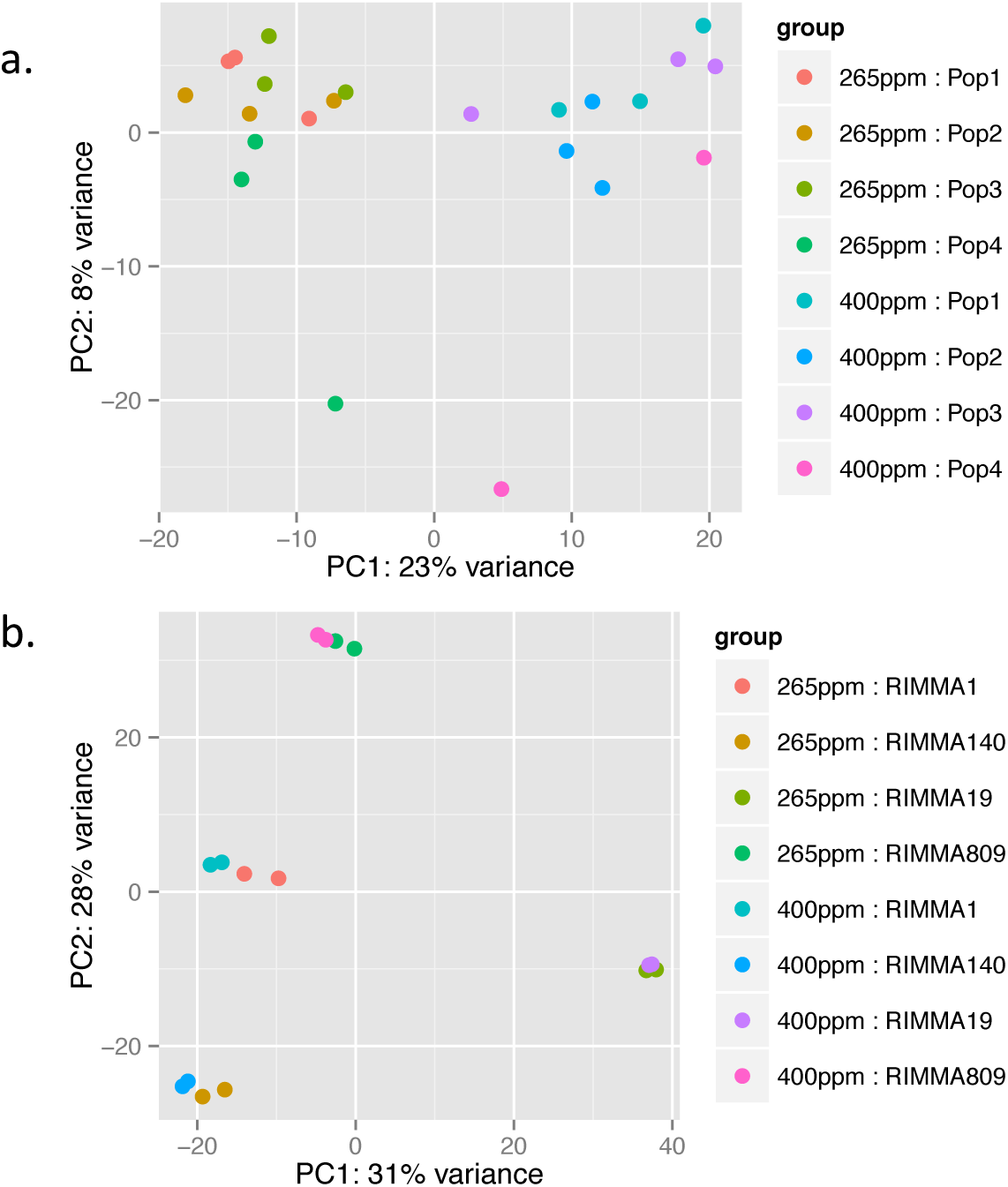
Principal Component Analysis (PCA) using rlog-normalized of the expression data for the principal components 1 (PC1) and PC2.

**Figure 3.**
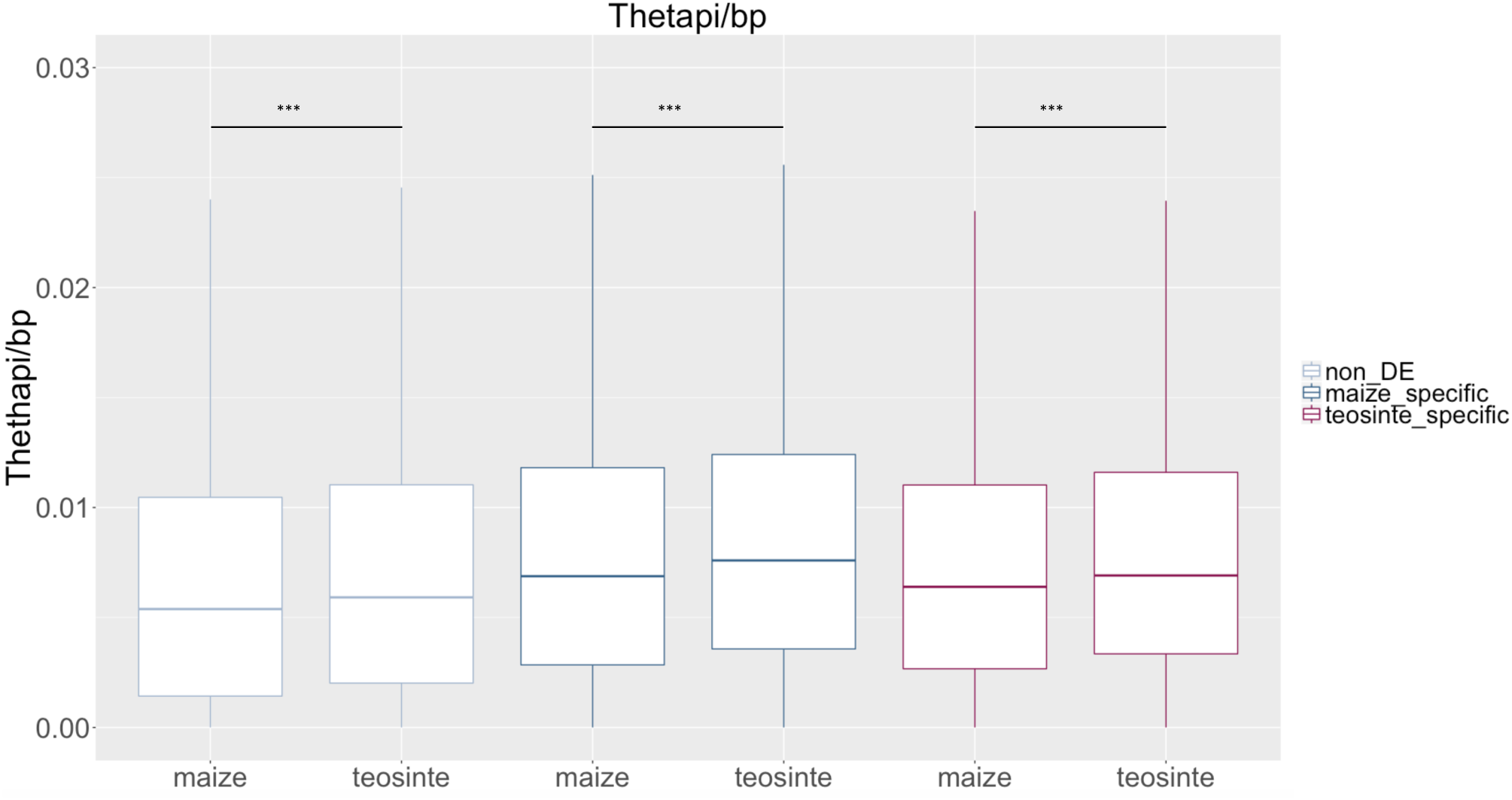
Nucleotide diversity. calculated for modern maize inbred lines and teosintes for the non-DE genes in gray, maize-specific genes in blue and the teosinte-specific DE genes in red. Mann-Whitney U tests for all comparisons are significant (***P <0.001)

